# Nitrate inhibition of nodule formation in *Medicago truncatula* is mediated by *ACC SYNTHASE 10*

**DOI:** 10.1101/434829

**Authors:** Arjan van Zeijl, Kerstin Guhl, Ting Ting Xiao, Defeng Shen, René Geurts, Wouter Kohlen

## Abstract

Legumes form a mutualistic endosymbiosis with nitrogen-fixing rhizobia. These rhizobia are housed intracellularly in specialised lateral root organs, called nodules. Initiation of these nodules is triggered by bacterial derived signalling molecules, lipochitooligosaccharides (LCO). The process of nitrogen fixation is highly energy-demanding and therefore nodule initiation is tightly regulated. Nitrate is a potent inhibitor of nodulation. However, the precise mechanisms by which nitrate inhibits nodulation is poorly understood. Here, we demonstrate that in *Medicago truncatula* nitrate interferes with the transcriptional regulation of the ethylene biosynthesis gene *ACC SYNTHASE 10*. ACSs commit the rate limiting step in ethylene biosynthesis and in *M. truncatula ACS10* is highly expressed in the zone of the root where nodulation occurs. Our results show that a reduction in *ACS10* expression in response to LCO exposure correlates with the ability to form nodules. In addition, RNAi-mediated knockdown of *ACS10* confers nodulation ability under otherwise inhibitory nitrate conditions. This discovery sheds new light on how ethylene is involved in the inhibition of nodulation by nitrate, bringing us one step closer to understanding how plants regulate their susceptibility towards rhizobia.

## INTRODUCTION

Legumes can engage in a nitrogen-fixing endosymbiosis with diverse bacterial species collectively called rhizobia (for review see Ferguson et al., 2018). In this interaction, the rhizobium bacteria are accommodated intracellularly in specialized root organs called nodules. These nodules create optimal physiological conditions for the conversion of atmospheric nitrogen into ammonium using the bacterial enzyme nitrogenase. This process is energetically costly and therefore nodule formation is strictly regulated. Understanding how legumes regulate nodule formation based on internal and external cues could provide tools to expand the agricultural application of rhizobial nitrogen-fixation (Mus et al., 2016).

The legume-rhizobium symbiosis is initiated upon perception of rhizobial-secreted lipochitooligosaccharide (LCO) signals (for review see Limpens et al., 2015). Perception of these signals mitotically activates cells in the root cortex and pericycle, leading to nodule primordium formation. In addition, LCO perception also initiates an infection process that starts in the root hair cell that perceives the LCO signal. This involves a redirection of growth, resulting in the formation of a tight curl that entraps the bacteria in an infection pocket. From there, cell wall-bound infection threads are formed that guide the bacteria to the developing nodule primordium. Once inside the nodule, the rhizobia are released into the host cell cytoplasm as transient nitrogen-fixing organelles. These so-called symbiosomes remain surrounded by a host-derived membrane, which facilitates exchange of ammonium and photo-assimilates between host and microbe.

Rhizobium LCOs are perceived by a specific set of receptors at the root epidermis. This activates a signalling cascade that results in nuclear calcium oscillations. These oscillations are decoded by a nuclear-localized Ca^2+^/calmodulin-dependent protein kinase (CCaMK). CCaMK phosphorylates the transcription factor CYCLOPS, resulting in massive transcriptional reprogramming (Breakspear et al., 2014; Larrainzar et al., 2015; Van Zeijl et al., 2015). Among the transcriptional targets of CYCLOPS is *NODULE INCEPTION* (*NIN*), a master regulator of symbiotic signalling (Singh et al., 2014). Activation of CCaMK also triggers an accumulation of the bio-active cytokinins isopentenyl adenine (iP) and *trans*-zeatin (tZ) (Van Zeijl et al., 2015). Mutants affected in cytokinin perception develop severely reduced nodule numbers, indicating that activation of cytokinin signalling is required for nodule formation (Murray et al., 2007; Plet et al., 2011; Held et al., 2014)

In contrast to the positive role of cytokinin on nodule formation, ethylene negatively regulates rhizobial infection and nodule initiation (Oldroyd et al., 2001). Ethylene is produced through consecutive activity of 1-aminocyclopropane-1-carboxylic acid (ACC) synthases (ACS) and ACC oxidases (ACO) (Kende, 1993). The conversion of *S*-adenosylmethionine into ACC through ACS activity is generally considered rate limiting (Kende, 1993). Several ACS-encoding genes in *Medicago truncatula* are transcriptionally induced in response to LCO signalling (Larrainzar et al., 2015; Van Zeijl et al., 2015; Herrbach et al., 2017), and an increase in ethylene release has been reported for *Lotus japonicus* at six hours after inoculation (Reid et al., 2018). The *sickle* (*skl*) mutant in *M. truncatula* is ethylene insensitive due to a mutation in *EIN2*, which encodes a central component in the ethylene signalling cascade (Penmetsa and Cook, 1997; Penmetsa et al., 2008). The *skl* mutant is hyper-infected by rhizobium and forms a multitude of nodules in distinct clusters along the root (Penmetsa and Cook, 1997). A similar phenotype is observed in *Lotus japonicus*, when both EIN2-encoding genes are mutated (Miyata et al., 2013; Reid et al., 2018), indicating an important role for ethylene signalling in the regulation of nodulation.

In most legumes, rhizobial susceptibility is negatively regulated by exogenous sources of fixed nitrogen (Carroll and Gresshoff, 1983; Streeter, 1985; Wong, 1988). The effect of fixed nitrogen on rhizobial susceptibility appears gradual and depends on the concentration of nitrogen that is applied to the plant. At low concentrations of nitrogen, nitrogen fixation rates are decreased, whereas at higher concentrations rhizobial infection and nodule formation could be completely abolished (Streeter, 1985). Several studies have investigated the inhibition of nodulation by fixed nitrogen sources. However, the data appears scattered and the effects of nitrogen on nodulation differ between legume species and sources of nitrogen (ammonium vs. nitrates). For example, in vetch (*Vicia sativa*) and *L. japonicus*, application of 10 mM of NH_4_NO_3_ blocks rhizobial infection already at the stage of root hair curling, an effect not observed after application of KNO_3_ (Heidstra et al., 1994; Barbulova et al., 2007).

It has been demonstrated that high concentrations of KNO_3_ inhibit the induction of *NIN* in *L. japonicus* at 24 hours after rhizobial inoculation or LCO application, an effect that is not observed in a *har1* mutant (Barbulova et al., 2007). The *L. japonicus har1* mutant is affected in autoregulation of nodulation (AON), a mechanism that restricts nodule numbers based on signals released from already existing nodules (Ferguson et al., 2010; Reid et al., 2011; Gresshoff and Ferguson, 2017). Mutants in this pathway are super-nodulated and appear partially resistant to nitrogen inhibition of nodulation (Barbulova et al., 2007). In addition, partial resistance to inhibition of nodulation by nitrogen is also observed in alfalfa (*Medicago sativa*) plants treated with the ethylene synthesis inhibitor aminoethoxyvinylglycine (Ligero et al., 1991). Furthermore, the release of ethylene from inoculated alfalfa roots is increased after addition of nitrates (Ligero et al., 1987). Combined this suggests that both AON and ethylene function as signalling intermediates during inhibition of nodulation by fixed forms of nitrogen. However, despite these insights exactly how and at which point during nodule formation inhibition by nitrogen occurs remains unclear.

Here, we set out to investigate the mechanisms by which nitrate inhibits nodulation in *M. truncatula*. We demonstrate that a plate system can be used to study the effect of nitrate on nodulation in *M. truncatula*. We use this system to examine at which point during nodule initiation inhibition by nitrate occurs and come to a model how ethylene contributes to this inhibition.

## RESULTS

The effect of nitrate on nodulation has been reported before. However, results are often fragmented (Carroll and Gresshoff, 1983; Streeter, 1985; Wong, 1988). In an attempt to investigate the mechanism by which nitrate inhibits nodule initiation in *Medicago truncatula* we turned to an agar-solidified Fåhraeus plate system. For this, seedlings of *M. truncatula* were grown on medium with increasing concentrations of Ca(NO_3_)_2_. This results in nitrate concentrations ranging from 0 to 16 mM. The concurrent increase in Ca_2_^+^ concentration was compensated by addition of CaCl_2_ (Fig 1a, Supplemental table S1). Roots were grown in semi-darkness to prevent ethylene accumulation, which allows efficient nodule formation without addition of the ethylene synthesis inhibitor 2-aminoethoxyvinyl glycine (AVG) (Supplemental figure S1).

**Figure 1.**
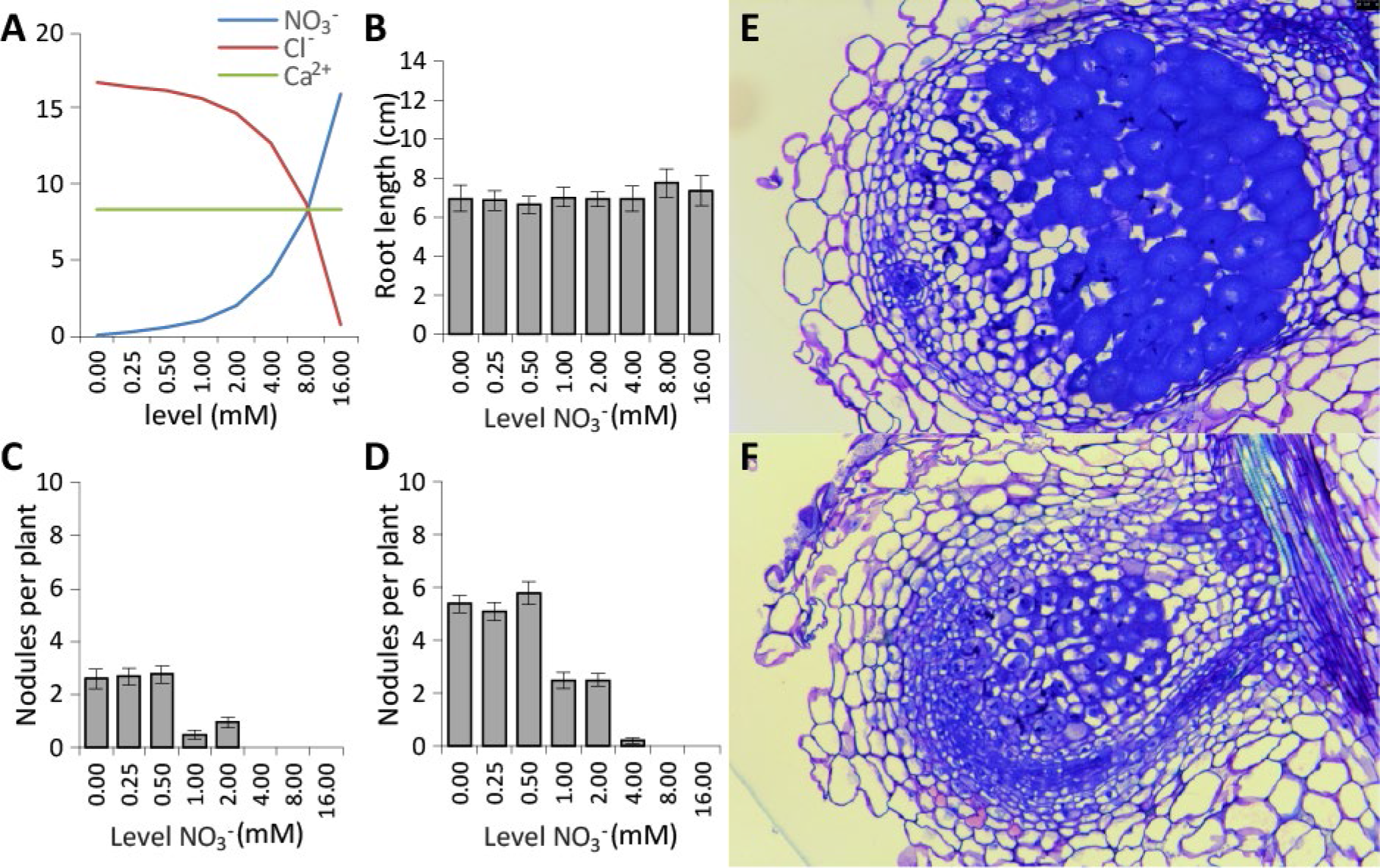
Nitrate inhibition of nodulation on Fåhraeus plates. **(A)** Schematic representation of the increase in nitrate levels, **(B)** Effect of increasing nitrate concentrations on primary root growth at 7 DPI (days post inoculation), **(C)** Nodule numbers at 7 DPI, **(D)** Nodule numbers at 14 DPI, **(E)** Section of a nodule formed on 0.25 mM NO_3_^-^, **(F)** Section of a nodule formed on 2 mM NO_3_^-^, (for all n=40).

After 5 days of growth, plants were inoculated with *Sinorhizobium meliloti* strain 2011 at which point root length was determined. No effect of nitrate was observed (Fig 1b), indicating that the varying concentrations of nitrate have no effect on root development. Nodule numbers were scored at 7 days post inoculation (DPI) and 14 DPI (Fig 1c-d). At 7 DPI, nodules developed normally on plants grown on 0, 0.25 and 0.5 mM NO_3_^-^. On plants grown at 1 and 2 mM NO_3_^-^ nodule numbers were reduced, and on plants grown at 4, 8 and 16 mM NO_3_^-^ nodules were not observed at this time point (Fig 1c). A similar picture emerged at 14 DPI. Average nodule numbers dropped by 50% when nitrate concentrations in the medium increased from 0.5 to 1 mM. In contrast to 7 DPI, a few nodules were observed on plants grown at 4 mM NO_3_^-^, but not on plants grown at 8 or 16 mM NO_3_^-^ (Fig 1d) In addition to reduced nodule numbers, nodules formed on 1 and 2 mM NO_3_^-^ remained white, suggesting that these nodules are unable to fix nitrogen. Sectioning revealed that nodules formed on 0.25 mM NO_3_^-^ developed normally and display a zonation characteristic of indeterminate nodules (meristem, infection, fixation zone, Fig 1e). Nodules formed on 2 mM NO_3_^-^ also developed a meristem and infection zone. However, a stable fixation zone was not observed. Instead, infected cells seemed to senesce almost instantly (Fig 1f). To determine at which point during nodule initiation interference by nitrate occurs in our system, we examined whether root hair deformation, and initiation of cell division could be observed. To analyse root hair deformation, seedlings were transferred to Fåhraeus slides 24 hours after germination and grown for an additional 2 days on different concentrations of nitrate. Treatment with ^~^10^-9^ M Sm2011 LCOs for 3 h induced root hair curling on all tested nitrate concentrations with no apparent differences between concentrations, indicating that root hair curling in our setup is not susceptible to inhibition by nitrate (Fig 2a-i)

**Figure 2.**
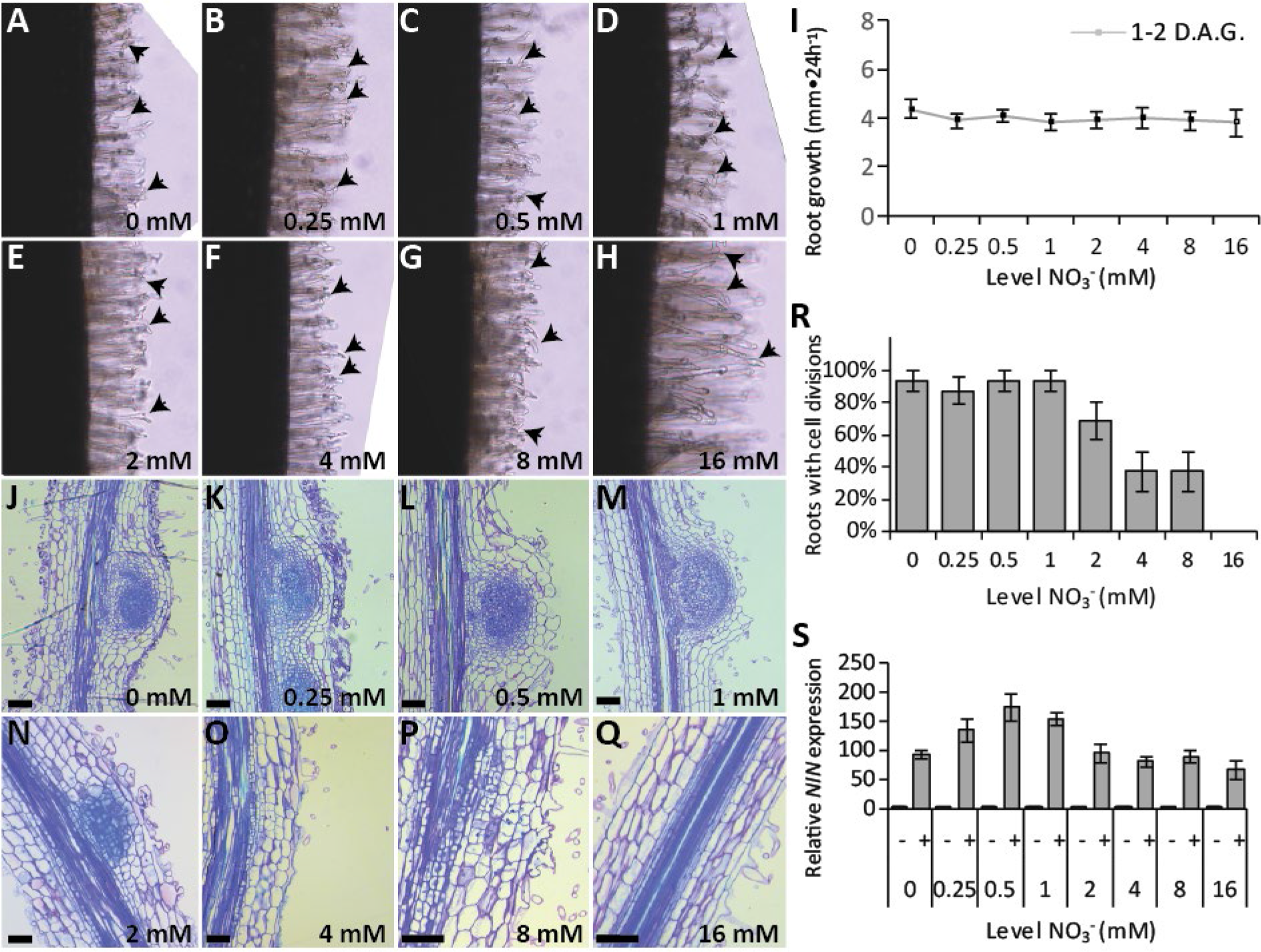
The effect of increased nitrate levels on early nodulation responses in wild-type *Medicago truncatula*. **(A-H)** Root hair deformations at increasing levels of nitrate (arrows point at curled roots). Pictures are representative of 10 biological replicates, **(I)** The effect of increasing levels of nitrate on primary root growth at 2 DAG (days after germination), **(J-Q)** Sections through the root susceptible zone at 4 DPI. Pictures are representative of 10 biological replicates, **(R)** Percentage of roots showing any cell divisions under increasing levels of nitrate at 4 DPI (n=10), **(S)** The effect of nitrate on LCO induced *NIN* expression (n=3).

Next, we determined at which nitrate concentrations cell division could be initiated following rhizobial inoculation. To this end, the root susceptible zone of plate-grown *M. truncatula* was inoculated with 10 μL *S. meliloti* (OD_600_ = 0.1) and sectioned at 4 DPI. This showed that cell divisions could be initiated at all tested concentrations, except at 16 mM (Fig 2j-q). Although cell divisions were initiated at 4 and 8 mM NO_3_^-^, well-developed nodule primordia could not be observed (Fig 2o-q). In addition, the percentage of roots that initiate cell division at 4 and 8 mM NO_3_^-^ dropped roughly 50% compared to the lower concentrations (Fig 2r). Altogether, this suggests that in our system the ability to induce cell divisions is gradually reduced by increasing levels of nitrate.

Next, we tested whether early induction of LCO signalling genes is affected by nitrate. For this, we focused on *NODULE INCEPTION* (*NIN*). Under control conditions, *NIN* expression is up-regulated within 3 hours of LCO application (Van Zeijl et al., 2015). Interestingly, *NIN* expression was induced across the entire nitrate concentration range (Fig 2s). However, the strongest induction of *NIN* was observed at the lower nitrate concentrations (i.e. 0.25-1 mM).

It has been demonstrated that LCO signalling triggers a rapid accumulation of cytokinins in the *M. truncatula* root susceptible zone (Van Zeijl et al., 2015). To determine whether this response is affected by nitrate, we measured cytokinin concentrations at 3 hours after LCO exposure. The level of *cis*-zeatin (cZ) and *cis*-zeatin riboside (cZR) was unaffected by either LCO application or nitrate concentrations (Fig 3a-b). At low nitrate concentrations (0-0.5 mM), LCO treatment induces a strong accumulation of the bioactive cytokinins isopentenyl adenine (iP) and *trans*-zeatin (tZ) as well as their riboside-derivatives iPR and tZR (Fig 3c-f). At higher concentrations of nitrate, this response is gradually reduced. At 0.25 mM NO_3_^-^, iP and tZ levels increase by approximately 3- and 12-fold, respectively. In contrast, at 16 mM NO_3_^-^ iP and tZ concentrations increase by only 1.5- and 2.5-fold, respectively. This indicates that NO^-^_3_-at least in part-blocks LCO induced accumulation of cytokinin in the Medicago root susceptible zone.

**Figure 3.**
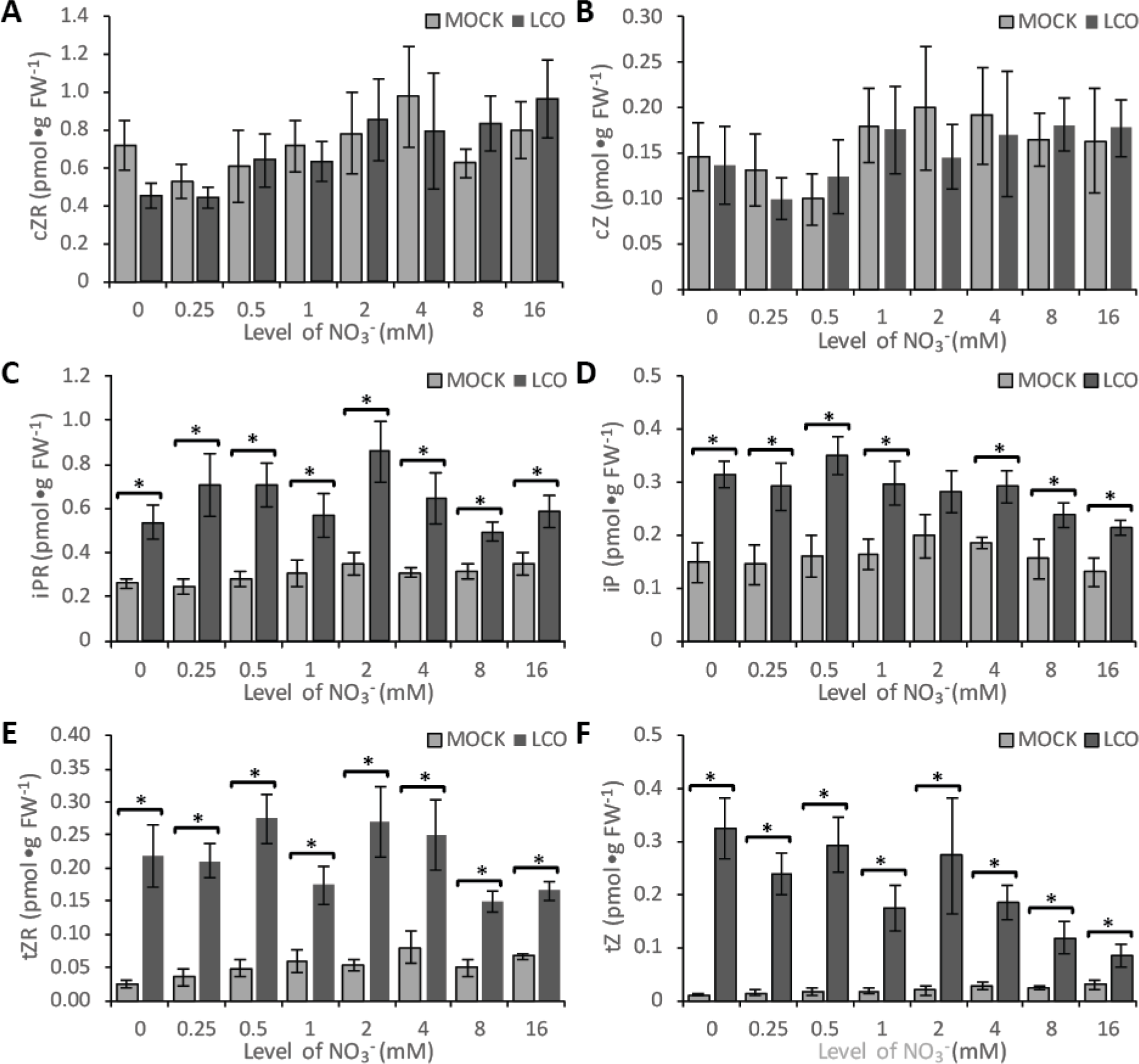
The effect of increasing nitrate levels on cytokinin accumulation in the root susceptible zone of wild-type *Medicago truncatula*. The levels of; **(A)** cZR, **(B)** cZ, **(C)** iPR, **(D)** iP, **(E)** tZR, and **(F)** tZ per gram fresh weight measured by UPLC-MS/MS. * = statistical difference (p<0.05) between mock and LCO application (student t-test, n=6).

### Inhibition of nodulation by nitrate is mediated by ethylene

Ethylene negatively affects LCO signalling and has been postulated to be involved in the inhibition of nodulation by nitrate (Ligero et al., 1991; Lee and LaRue, 1992; Caba et al., 1998). To determine whether the gradual reduction of symbiotic responses by nitrate in our system is mediated by ethylene, we determined nodulation behaviour of the ethylene-insensitive *sickle* mutant. Consistent with previous reports, the *sickle* mutant forms about 8 times more nodules than wild type at low nitreate concentrations (Fig 4a; (Penmetsa and Cook, 1997)). At 2 mM NO_3_^-^, average nodule number is reduced by 50% on *sickle* mutant roots (Fig 4a-c). Interestingly, increasing concentrations of nitrate do not cause a further reduction in nodule numbers (Fig 4a). Instead, a comparable number of nodules was formed on *sickle* plants grown at nitrate concentrations ranging from 2 to 16 mM (Fig 4a). Consistently, cytokinin measurements after LCO exposure indicate a strong accumulation of iP and tZ in roots of the *sickle* mutant grown at 0.25 or 16 mM NO_3_^-^ (Fig 4d-g). At 0.25 mM NO_3_^-^, iP and tZ concentrations increase roughly 12 and 100 times after LCO treatment. At 16 mM NO_3_^-^, this response is reduced by 50%, but still exceeds the response in wild-type roots grown at 0.25 mM NO_3_^-^ (Fig 4d-g). Together, this indicates that ethylene-insensitivity creates a partial resistance to the inhibition of nodulation by nitrate.

**Figure 4.**
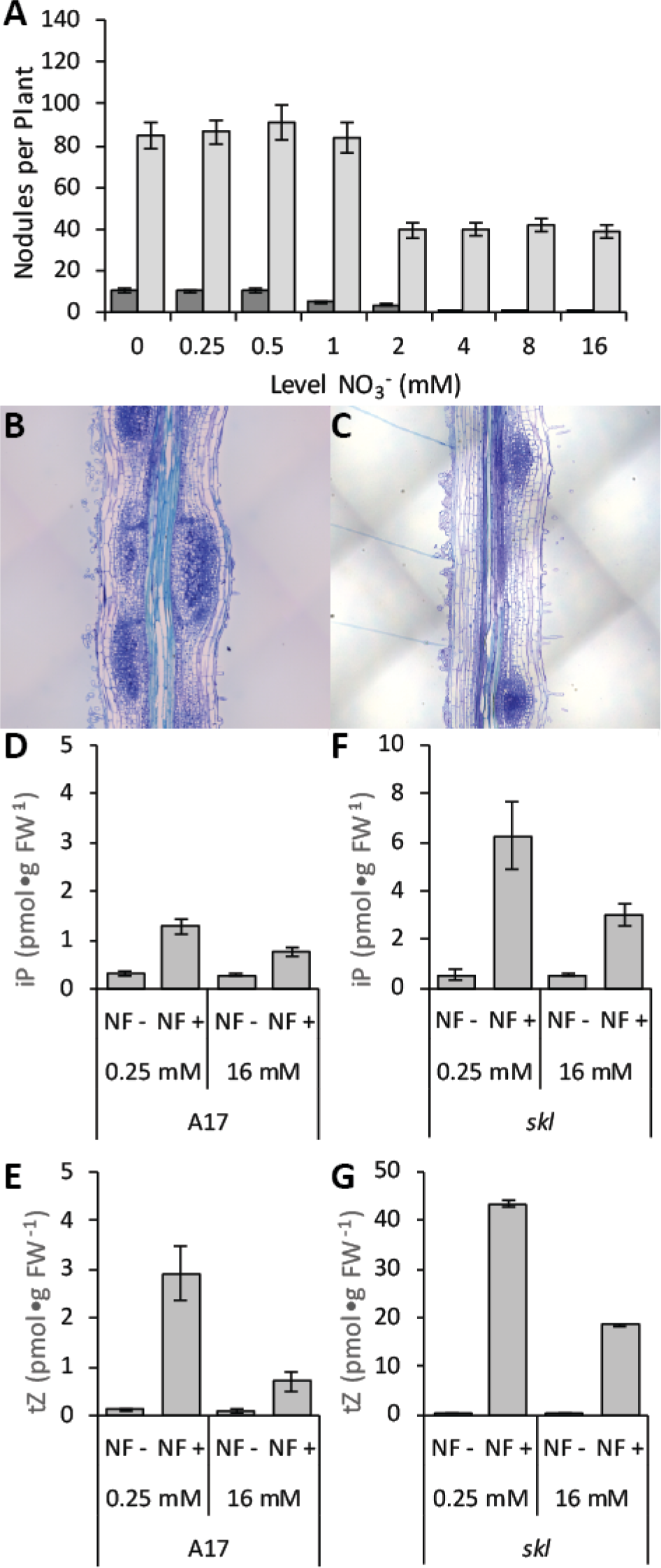
The effect of increased nitrate levels on nodulation in the *Medicago truncatula sickle/ein2* mutant. **(A)** Effect of increased nitrate on nodulation at 7 DPI (days post inoculation, n=40). Sections through the root susceptible zone of **(B)** *Mtsickle* and **(C)** wild-type *M. truncatula* at 4 DPI. Pictures are representative of 10 biological replicates. The levels of; iP and tZ in **(D, E)** wild type and **(F, G)** *Mtsickle* per gram fresh weight measured by UPLC-MS/MS (n=6).

To determine how ethylene is involved in the inhibition of nodulation by nitrate, we determined expression of ethylene biosynthesis genes. None of the ACC oxidase-encoding genes was responsive to nitrate or LCO treatment (Fig 5a). Similarly, expression of an ACC deaminase was not affected by either treatment (Fig 5b). Ten ACC synthases (ACS) are encoded in the *M. truncatula* genome of which only 4 are LCO-responsive (Fig 5c-f). *ACS1, ACS2* and *ACS3* are induced by LCOs (Van Zeijl et al., 2015), However in these experiments only two, *ACS1* and *ACS3* were detected (Fig 5c). Both were similarly induced at lower and higher nitrate levels (Fig 5d, e, Supplemental figure S2a, b). *ACS10* is highly expressed in roots of *M. truncatula* (Fig 5f). At 0.25 mM NO_3_^-^, expression of *ACS10* is reduced by 50% in response to LCO treatment, a response that is not observed in roots grown at 16 mM NO_3_^-^ (Fig 5f, Supplemental figure S2c).

**Figure 5.**
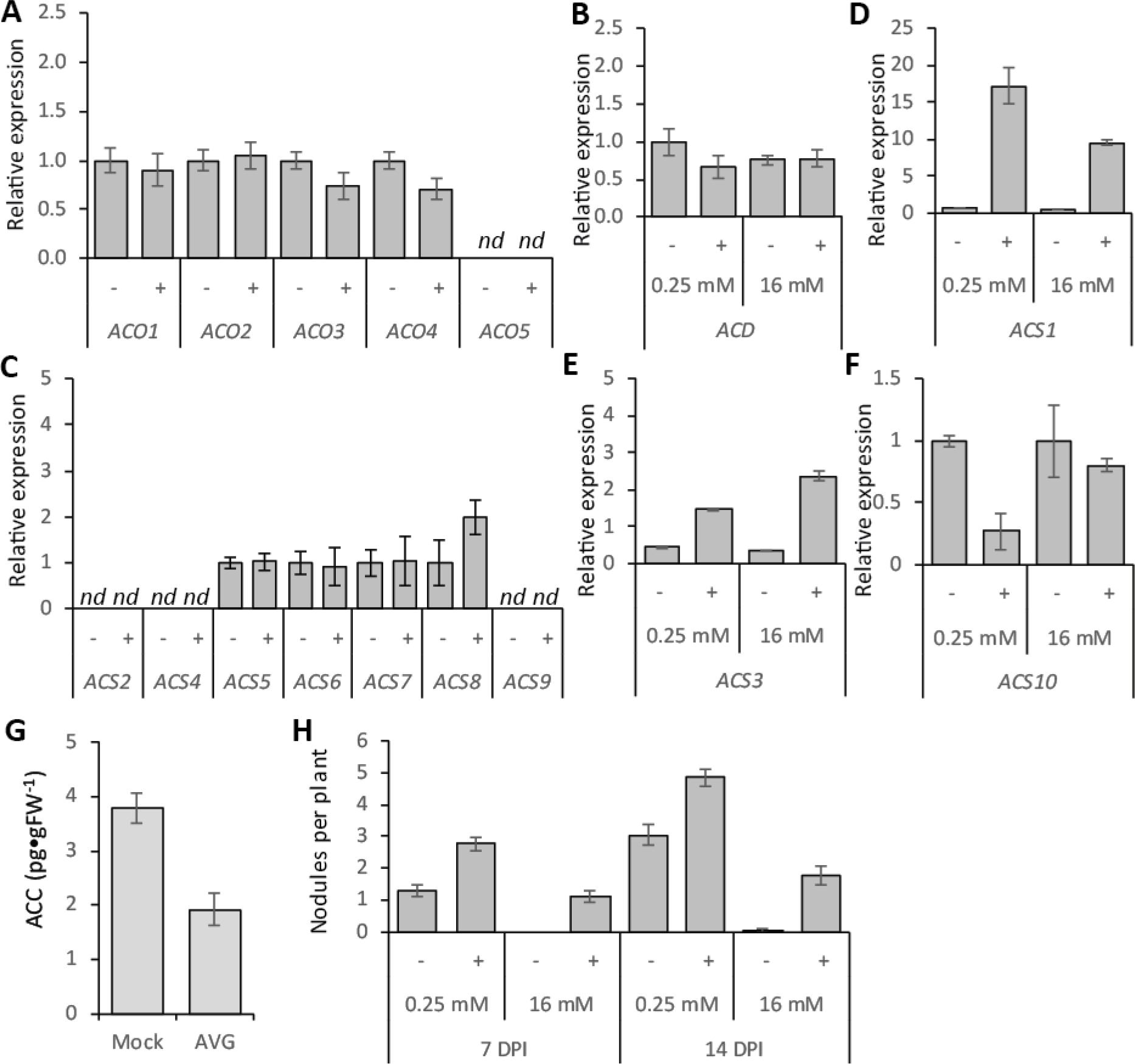
Analysis of ethylene biosynthesis. Effect of mock (−) and LCO signalling (+) on **(A)** *ACO1-5*, **(B)** *ACS2, 4-9*. Effect of mock (−) and LCO signalling (+) under low and high nitrate levels on **(C)** *ACD*, **(D)** *ACS1*, **(E)** *ACS3*, and **(F)** *ACS10*. **(G)** The effect of AVG application on ACC levels after 7 days of exposure. **(H)** Effect of AVG application on nitrate inhibition during nodulation. All gene expression analysis were performed at 3h post inoculation (n=3). AVG application, measurements n=6, nodulation n=40.

To determine whether a reduction in ACS activity confers resistance to nitrate in *M. truncatula*, we grew seedlings on medium containing 2 μM AVG. This reduced ACC concentrations in the root susceptible zone by ^~^50% (Fig 5g). At 0.25 mM NO_3_^-^, AVG treatment increased nodule numbers by 30-50%. At 16 mM NO_3_^-^, nodules were formed on plants treated with AVG, but not on control plants (Fig 5h). This indicates that a reduction in ACC content creates a partial resistance to nitrate.

Next, we determined whether a reduction in *ACS10* expression also confers resistance to inhibition of nodulation by nitrate. To this end, we created compound plants bearing transgenic roots in which *ACS10* expression was reduced by RNAi. RNA-mediated silencing specifically reduced *ACS10* expression by 75% (Fig 6a, Supplemental figure S3), resulting in a 35% decrease in ACC concentration (Fig 6b). At 0.25 mM NO_3_^-^, roots expressing the RNAi construct against *ACS10* formed 2 times more nodules than non-transformed roots or roots expressing a control construct (Fig 6c). At 16 mM NO_3_^-^, nodules were only formed on roots expressing the RNAi construct against *ACS10*. Nodule numbers on these roots were comparable to those on non-transgenic roots or roots transformed with a control construct grown at 0.25 mM NO_3_^-^ (Fig 6d). This indicates that silencing of *ACS10* creates resistance to the inhibition of nodulation by nitrate. This effect is specific to *ACS10*, as silencing of *ACS1* and *ACS3* together does not confer resistance to nitrate (Supplemental figure S4). This despite the fact that expression of *ACS3* alone is equal to the reduction in *ACS10* expression observed after LCO treatment at 0.25 mM NO_3_^-^ (Fig 5e, f). Taken together, this suggest that nitrate blocks nodule formation by inhibiting LCO-mediated repression of *ACS10* expression.

**Figure 6.**
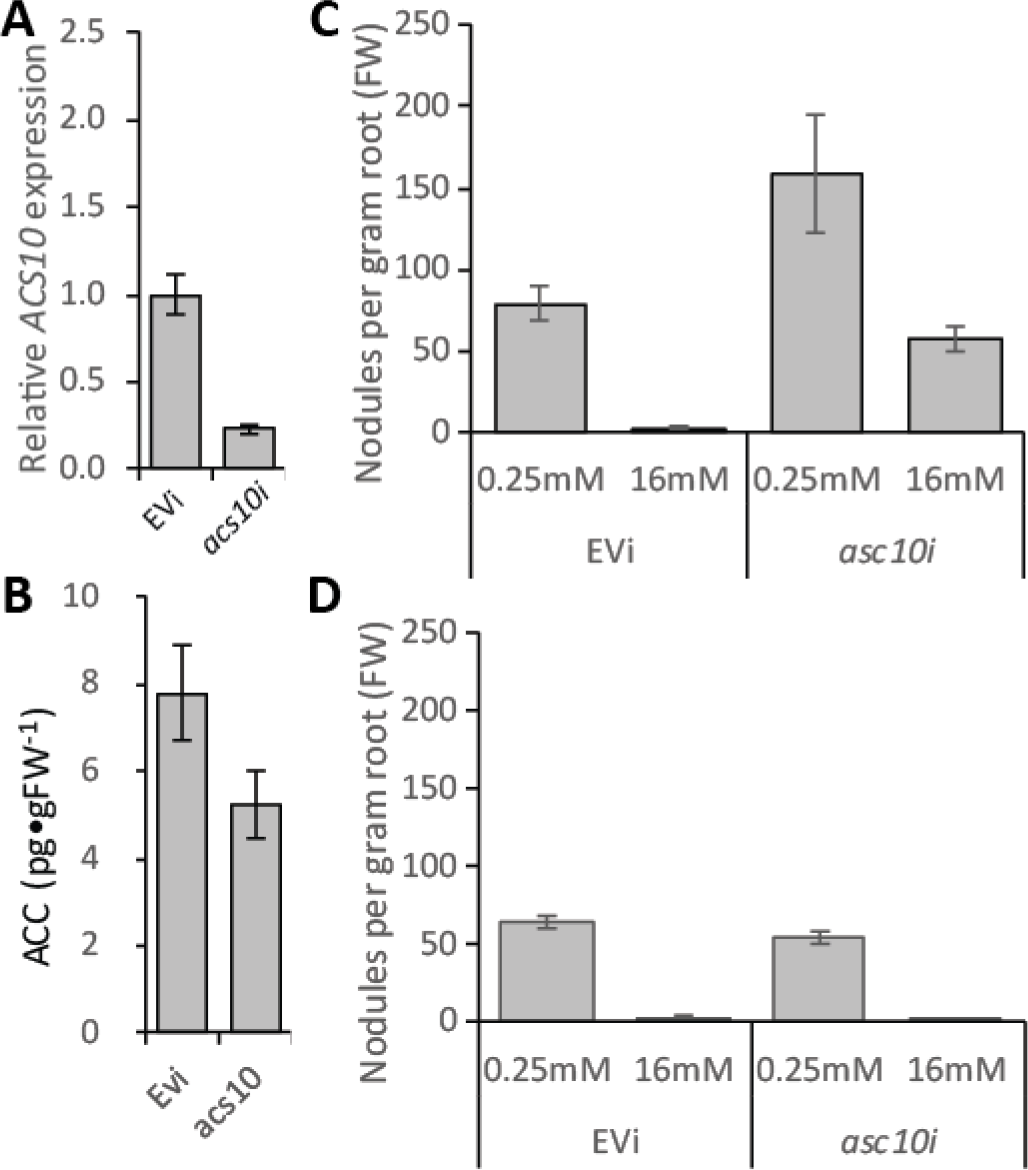
The effect of RNAi-mediated *ACS10* knockdown on nodulation. **(A)** Relative expression of *ACS10* in the empty vector control (*EVi*) and *ACS10* RNAi targeted roots (*ACS10i*) (n=3). **(B)** Effect of *ACS10* silencing on the ACC levels in the root susceptible zone (n=6). The effect of low and high nitrate on nodule formation on **(C)** roots transformed with the *EVi* and *ACS10i* constructs and **(D)** the untransformed roots formed on the same compound plants (n=10).

## DISCUSSION

Here, we aimed to investigate the inhibition of nodulation by nitrate in *Medicago truncatula*. For this, we used a plate system in which nodule formation is completely blocked at 4 mM NO_3_^-^. We show that this inhibition of nodulation by nitrate can be partially overcome through interference with ethylene biosynthesis or signalling. Furthermore, we show that the inhibition of nodulation by nitrate is mediated by *ACS10*. *ACS10* is highly expressed in *M. truncatula* roots and its expression is reduced by >50% after LCO recognition, a response that is not observed in roots treated with nitrate. RNAi-mediated silencing of *ACS10* confers resistance to inhibition of nodulation by nitrate. This strongly suggests that a reduction in *ACS10* expression in *M. truncatula* roots following LCO recognition is required for the initiation of cell divisions associated with the formation of nitrogen-fixing nodules.

We developed a plate system that allows efficient nodulation without the addition of chemical inhibitors (e.g. AVG). In this system, roots are shielded from light exposure by wrapping the bottom part of the plates in aluminium foil. The fact that both treatments, AVG application or light shielding, allow efficient nodulation suggests that the difficulties in nodulating *M. truncatula* on plates might be contributed to light-induced ethylene production.

In our plate system, the inhibitory effect of nitrate on nodulation seems to be a two-step process. Intermediate concentrations reduce nodule numbers by 50%. However, the nodules that do form do not contain a well-defined fixation zone and appear to be white, suggesting that these nodules are non-fixing. At higher concentrations of nitrate, nodules are only occasionally observed. This dual effect of nitrate is consistent with earlier reports that also described a higher sensitivity to nitrate with regard to nitrogen-fixation rates compared to nodule formation (Streeter, 1985). A possible explanation for this dual effect could be that in soil nitrogen concentrations are not uniformly distributed. Therefore, it would be unfavourable to block nodulation completely when roots pass through a nitrogen-rich patch. This would allow nodulation to continue almost instantly when nitrate levels drop below a certain threshold, if nodule initiation is not completely abolished.

We determine at which stage of nodule initiation inhibition by nitrate occurs. This showed that in our system nodule formation is blocked at 4 mM NO_3_^-^, whereas cell divisions associated with nodule primordium formation could still be observed at 8 mM NO_3_^-^. At 16 mM NO_3_^-^, cell divisions are fully inhibited while root hair deformation and NIN expression are not affected. Previous reports showed that root hair deformation is affected by ammonium nitrate treatment, but not by application of nitrate alone (Heidstra et al., 1994), consistent with our observations. In contrast to what we found, experiments in *L. japonicus* previously showed that NIN induction is affected by application of 10 mM NO_3_^-^ (Barbulova et al., 2007). The reason for this discrepancy is unclear. However, it could be related to the time point at which *NIN* expression is determined. In our experiments, *NIN* expression was determined at 3h after LCO exposure, a time point that is preceding initiation of cell division (Xiao et al., 2014). In the *L. japonicus* experiments, *NIN* expression was determined after 24h of LCO treatment or rhizobial inoculation. It is conceivable that at that time point cell divisions associated with nodule primordium formation have already been initiated. We show that application of 16 mM NO_3_^-^ completely blocks initiation of symbiotic cell divisions in *M. truncatula* roots. Therefore, it is possible that the reduction in *NIN* expression in *L. japonicus* in response to nitrate application is resulting from an inhibition of nodule initiation by nitrate, rather than a direct effect of nitrate on the induction of *NIN* by LCO signalling. However, in order to determine whether this might indeed be true, *NIN* expression would have to be determined at several time points after rhizobial inoculation in the presence of nitrate.

It has been shown that the induction of cell divisions associated with nodule primordium formation is a cytokinin–mediated process (Murray et al., 2007; Plet et al., 2011; Van Zeijl et al., 2015). Therefore, we wanted to know whether nitrate inhibition is interfering with the cytokinin-mediated initiation of cell division. We showed that nitrate treatment did not affect the basal cytokinin levels at any of the tested concentrations. Surprisingly, LCO application induced accumulation of cytokinin within 3h even at high levels of nitrate. However, there is a clear decrease in the magnitude of this response at the higher nitrate levels. This reduced response correlates to some extent with the observed reduction in symbiotic cell divisions at increasing nitrate concentrations. However, despite a small increase in cytokinin concentration at 16 mM NO_3_^-^, cell divisions were not observed at this concentration. Combined, this suggests that the inhibition of cell divisions is unlikely to be related to cytokinin levels alone.

Ethylene has been shown to negatively regulate nodule formation. Additionally, previous studies suggested a role for ethylene in the regulation of nodulation by nitrate (Ligero et al., 1987; Ligero et al., 1991; Caba et al., 1998). Application of AVG was shown to create partial resistance to nitrate inhibition of nodulation in alfalfa (Ligero et al., 1991). We show that in *M. truncatula*, AVG application reduced ACC concentrations in roots by ^~^50%, which is sufficient to allow nodule formation at 16 mM NO_3_^-^. Consistently, the ethylene insensitive *skl* mutant forms a large number of nodules at all tested nitrate concentrations. However, this mutant is not fully resistant to nitrate inhibition of nodulation. A two-fold reduction in nodule numbers is still observed in the *skl* mutant between 1 and 2 mM NO_3_^-^. Similarly, the accumulation of cytokinin following LCO application at 16 mM is ^~^50 reduced compared to that at 0.25 mM. Together this suggests that ethylene is not the only factor involved in nitrate inhibition of nodulation. Indeed, it has been shown that the AON pathway is also contributing to the inhibition of nodulation by nitrate (Barbulova et al., 2007; Ferguson et al., 2010; Reid et al., 2011). The two-fold reduction in nodule numbers in the *skl* mutant under elevated nitrate levels suggests that AON works in parallel to the ethylene regulation reported here. To test this hypothesis, analysis of a *skl sunn* double mutant (*SUNN* is the *M. truncatula* orthologue of *HAR1* in *L. japonicus* (Nishimura et al., 2002; Murray et al., 2006; Reid et al., 2011) could be performed. Such studies could also uncover whether besides AON and ethylene other regulatory pathways are involved in the inhibition of nodulation by nitrate.

To determine how ethylene is involved in the inhibition of nodulation by nitrate, we studied expression of ethylene biosynthesis genes. Research on *Arabidopsis thaliana* showed that nitrate induces the expression of these genes (Tian et al., 2009). However, we did not observe such regulation by nitrate in *M. truncatula*. Several ethylene biosynthesis genes have been shown to be induced by LCO application or rhizobial inoculation, including *ACS1* and *ACS3* (Breakspear et al., 2014; Larrainzar et al., 2015; Van Zeijl et al., 2015). This corresponds to an increase in ethylene release measured after rhizobial inoculation in *L. japonicus* (Reid et al., 2018). This response is likely part of a negative feedback loop to restrict rhizobial infections after a successful interaction has been initiated (Van Zeijl et al., 2015). Our data indicates that nitrate does not interfere with the induction of *ACS1* and *ACS3* by LCOs, suggesting that this putative negative feedback response is still functional. Instead, we found that nitrate interferes with the downregulation of *ACS10* expression following LCO application. *ACS10* is highly expressed in the *M. truncatula* root susceptible zone and its expression is reduced by ^~^50% following LCO exposure. This reduction in *ACS10* expression is gradually inhibited at increasing concentrations of nitrate. In this respect, the expression of *ACS10* is negatively correlated to the induction of cell divisions associated with nodule primordium formation. To determine whether this inhibition of transcriptional regulation of *ACS10* by LCOs is contributing to the inhibition of nodulation by nitrate, we reduced *ACS10* expression by RNAi. As such, *ACS10* expression was reduced by ^~^70%, which lead to a ^~^30% drop in ACC concentration. This reduction showed to be sufficient to allow nodule formation at 16 mM nitrate, a concentration that complete blocks nodule formation on control roots. A similar effect is not observed after RNAi of both *ACS1* and *ACS3*. Absolute expression of *ACS3* alone is equal to the reduction in *ACS10* expression induced by LCO signalling, indicating that resistance to nitrate inhibition of nodulation is a specific effect of *ACS10* knockdown and cannot be achieved by an overall reduction in *ACS* expression. Altogether, this strongly suggests that the inhibition of nodulation by nitrate involves interference with the transcriptional regulation of *ACS10* by LCOs. Additionally, it suggests that a reduction in *ACS10* expression is required for cells in the root cortex and pericycle to become competent to re-enter mitosis following recognition of rhizobial signals.

In conclusion, our data strongly indicates that inhibition of nodulation by nitrate involves interference with the transcriptional regulation of *ACS10* by LCO signalling. Additionally, these data suggest that a reduction in *ACS10* expression is a prerequisite for cells in the root susceptible zone to be competent to initiate cell divisions leading to nodule primordium formation. This finding could provide an important clue for transferring the nitrogen-fixing symbiosis to non-legume crops, a long-lasting dream in symbiosis research (Mus et al., 2016).

## MATERIALS AND METHODS

### Plant material, growth conditions and treatments

*Medicago truncatula* seeds (Jemalong A17 wild type, *Mtein2/Mtsickle* (Penmetsa and Cook, 1997) were treated with H2SO4 for 7 minutes, five times rinsed with MilliQ water and sterilized for 10 minutes using normal household-bleach. Seeds were washed with sterile MilliQ water for five times and placed on round Petri dishes containing Fåhraeus medium (0.25 mM NO_3_^-^, 1% Daishin agar) at 4°C in darkness for stratification. After 48h, seeds were transferred to RT to germinate for an additional 24h. Germinated seedlings were transferred to the appropriate *in vitro* growth system. For all experiments plants were grown in an environmentally-controlled growth chamber at 20°C/18°C with a 16h-light/8h-dark cycle and 70% relative humidity.

### Nitrate concentration range

A modified Fåhraeus medium (0.12 g/L MgSO_4_•7H_2_O, 0.10 g/L KH_2_PO_4_, 0.15 g/L Na_2_HPO_4_•2H_2_O, 1 ml/L 15 mM Fe-Citrate, 2.50 ml/L Spore-elements β- (CuSO_4_•5H_2_O 0.0354g/L, MnSO_4_•H_2_O 0.462g/L, ZnSO_4_•7H_2_O 0.974g/L, H_3_BO_3_ 1.269 g/L, Na_2_MoO_4_•2H_2_O 0.398 g/L), pH 6.7) (Fåhraeus, 1957) with incrementing levels of nitrate (0, 0.125, 0.25, 0.5, 1, 2, 4 and 8 mL 1M Ca(NO_3_)_2_) was used. Increasing Ca^2+^ levels were compensated by adding decreasing amounts (8.7, 8.575, 8.45, 8.2, 7.7, 6.7, 4.7 and 0.7 mL) of 1M CaCl_2_. For plate growth 1% Daishin agar was added. For root hair deformation, individual seedlings were transferred to Fåhraeus slides (Fåhraeus, 1957). Nodulation to square (12cm×12cm) plates with different nitrate concentrations. Eight seedlings were grown per plate and a minimum of 4 plates per replicate were grown. Plates were partially covered with tin foil to avoid light grown roots. All plants were grown under long day conditions (16 h of light 21 ⁰C and 8 h of darkness 18 ⁰C). From the growing root, susceptible zones (± 1 cm) of ca. 7-10 roots were collected and immediately frozen in liquid nitroge. Samples are stored for one day at -80 ⁰C.

### Nod-Factor or Rhizobia application

*Sinorhizobium meliloti* 2011 Nod factors (*Sm2011NF*) were purified and applied as previously described (Spaink et al., 1991; Van Zeijl et al., 2015) with minor modifications. *Sn2011NF* stocks were stored in 100% DMSO and diluted 100-fold (^~^10-9 M) in Fåhraeus medium. As mock treatment, Fåhraeus medium (1% DMSO) was used. Nitrate levels were kept equal to the growth condition of the seedlings. *Sn2011NF* were pipetted on top of the root susceptible zone. Roots were exposed for three hours and subsequently 1 cm root pieces were cut just above the root tip and snap-frozen in liquid nitrogen (n=6 for cytokinin measurements, n=3 for gene expression analysis). *Sinorhizobium meliloti* 2011 were grown and diluted to an OD_600_ of 0.1 or 0.01 for direct application on plates or inoculation in our semi sterile growth system, respectively.

### RNA isolation, cDNA synthesis and Quantitative RT-PCR

RNA was isolated from snap-frozen roots samples using the plant RNA kit (E.Z.N.A, Omega Biotek, Norcross, USA) according to the manufacturer’s protocol. From this, 1 μg total RNA was used to synthesize cDNA using the i-script cDNA synthesis kit (Bio-Rad, Hercules, USA) as described in the manufacturer’s protocol. Real time qRT-PCR was set up in 10 μl reactions with 2× iQ SYBR Green Super-mix (Bio-Rad, Hercules, USA). Experiments have been conducted on a CFX Connect optical cycler, according to the manufacturers protocol (Bio-Rad, Hercules, USA). All primers including the genes used for normalization (*MtUBQ10* and *MtPTB*) are given in Table SX. Data analysis was performed using CFX Manager 3.0 software (Bio-Rad, Hercules, USA). Cq values of 32 and higher were excluded from the analysis, though still checked for transcriptional induction (see Table SX). Statistical significance was determined based on student’s t-test (p<0.01).

### Vectors and constructs

For RNAi-mediated knockdown of *MtACS10* or *MtACS1*/*MtACS3*, two fragments (332-bp and 310-bp, respectively) were amplified from *M. truncatula* Jemalong A17 root cDNA, using specific primer pairs (Supplemental table S2), and cloned into pENTR-D-TOPO (Invitrogen, Carlsbad, USA). The both RNAi fragments were recombined into the DsRed-modified gateway vector pK7GWIWG2(II)-RR driven by the CaMV35S promoter (Limpens et al., 2005) to obtain the binary constructs pK7GWIWG2(II)-RR-p35S-MtACS10-RNAi and pK7GWIWG2(II)-RR-p35S-MtACS1/3-RNAi. For the empty vector control, the binary plasmid pK7GWIWG2(II)-RR-p35S-RNAi-control as previously described (van Zeijl et al., 2015) was used. All cloning vectors and constructs are available upon request from our laboratory or via the Functional Genomics unit of the Department of Plant Systems Biology (VIB-Ghent University).

### Plant transformation and histology

*Agrobacterium rhizogenes*-mediated root transformation was used to transform *M. truncatula* (Jemalong A17) as previously described (Limpens et al., 2004). Transgenic roots were selected based on *DsRED1* expression. Transgenic roots were transferred to low (0.125 mM Ca_2_(NO_3_)_2_) or high (4 mM Ca_2_(NO_3_)_2_) nitrate containing Fåhraeus with *Sinorhizobium meliloti* 2011 (OD_600_ = 0.01) roughly three weeks after transformation. After an additional three weeks of growth their nodulation phenotype was scored.

### Cytokinin and ACC extraction, detection and quantification by liquid chromatography-tandem mass spectrometry

Cytokinins analysis from the *M. truncatula* root were performed as previously described (Van Zeijl et al., 2015). ACC analysis from the *M. truncatula* root were performed as previously described (Bours et al., 2013).

## SUPPLEMENTARY DATA

**Supplemental figure S1.** Setting up the plate system for nodulation.

**Supplemental figure S2.** Analysis of the effect of nitrate on the LCO induction of the ethylene biosynthesis genes **(A)** *ACS1*, **(B)** *ACS3* and **(F)** *ACS10*. (n=3).

**Supplemental figure S3.** Off target effects of *ACS10i* and *ACS1-3i* on *ACS1*, *ACS2*, *ACS3*, *ACS4*, *ACS5*, *ACS6*, *ACS7*, *ACS8*, *ACS9*, and *ACS10*.

**Supplemental figure S4.** Nitrate inhibition on nodulation in pots, **(A)** The effect of 16 mM nitrate on nodulation in pots. The effect of ACS1-3i and ACS10i on 8 mM nitrate inhibition on nodulation in pots **(B)** untransformed roots, **(C)** transformed roots. **(D)** The effect of 8 mM nitrate on nodulation in pots.

**Supplemental table S1.** Sequences and primers used in this study.

## AUTHOR CONTRIBUTIONS

Conceptualization, A.v.Z. and W.K.; Methodology, K.G. and W.K.; Investigation, A. v. Z., K.G., T.T.X., D.S., and W.K.; Formal Analysis, A.v.Z., and W.K.; Visualization, W.K.; Writing – Original Draft, A.v.Z. and W.K.; Writing – Review & Editing, A.v.Z. and W.K.; Funding Acquisition, R.G. and W.K.; Supervision, W.K.

## ACKNOWLEDGMENTS

We would like to acknowledge Titis Wardhani, Rens Holmer, Jieyu Liu, Marijke Hartog, Sidney Linders, Henk Franssen and Ton Bisseling for their contributions to the project.

